# MDA5 and the integrated stress response control sex-specific type 1 diabetes onset in NOD mice

**DOI:** 10.64898/2026.07.02.736190

**Authors:** Erik Van Dis, Alessandra T. DeGidio, Lara Yao, Damion Winship, Carmela Sidrauski, Jacquelyn A. Gorman, Daniel B. Stetson

**Affiliations:** Department of Immunology, University of Washington School of Medicine, Seattle, WA 98109, USA; Calico Life Sciences LLC, South San Francisco, CA 94080, USA; Oklahoma Medical Research Foundation, Arthritis & Clinical Immunology, Oklahoma City, OK 73104, USA

## Abstract

Type I diabetes (T1D) is an autoimmune disorder in which the insulin producing cells of the pancreas are attacked and destroyed by autoreactive T cells. The innate immune mechanisms that contribute to T1D remain incompletely defined. Genome-wide association studies in humans have identified alleles of the *IFIH1* gene, which encodes the intracellular RNA sensor MDA5, that are strongly associated with development of T1D. We previously found that MDA5 signaling drives disease and mortality in a mouse model of Aicardi-Goutieres Syndrome (AGS) caused by mutations in the ADAR1 RNA editing enzyme. Genetic dissection of disease in this ADAR1 mutant mouse model revealed that the double stranded RNA-activated kinase PKR and the RNA sensor ZBP1 are also essential for disease. To test the role of intracellular RNA detection in T1D in the nonobese diabetic (NOD) mouse model, we used CRISPR targeting to generate NOD mice targeted for *Ifih1, Eif2ak2* (PKR) and *Zbp1*. We found that haploinsufficiency for *Ifih1* resulted in modest but significant protection from T1D only in male NOD mice, but neither PKR nor ZBP1 contributed to T1D onset or incidence. Moreover, treatment of NOD mice with a pharmacological inhibitor of the integrated stress response (ISR) had no effect on T1D incidence in female NOD mice, but accelerated and exacerbated disease in male NOD mice. Together, our findings demonstrate that MDA5 and the ISR contribute to sex-specific disease incidence in NOD mice.

## Introduction

Type 1 diabetes (T1D), in which the immune system attacks and destroys the insulin producing cells of the pancreas, remains a major clinical challenge. Despite decades of research into the genetics, origins, and progression of the disease, the only effective treatment for T1D remains injection of recombinant insulin combined with close monitoring of blood glucose levels^1^. Among mouse models of T1D, the non-obese diabetic (NOD) mouse has been studied the most extensively^2,3^. NOD mice develop spontaneous autoimmune diabetes, with faster onset and higher penetrance in females compared to males, similar to the human disease^2^. Immune cell infiltration into the pancreas precedes destruction of the insulin producing beta cells of the Islets of Langerhans, and both CD4 and CD8 T cells are essential for disease^2^. Despite the detailed understanding of the adaptive immune cells that drive diabetes progression, the innate immune signals that initiate and contribute to disease remain largely uncharacterized.

Genome-wide association studies have identified dozens of genomic loci that predict risk of T1D in humans^4,5^. Interestingly, the strongest single gene association with human T1D is the *IFIH1* gene that encodes MDA5, a RIG-I-like receptor that detects double stranded RNA (dsRNA) and initiates antiviral immunity through the production of type I interferons (IFNs)^6-8^. The MDA5 protein contains two N-terminal CARD domains that interact with the adapter protein MAVS to trigger antiviral signaling^9,10^, followed by a helicase domain that controls assembly of MDA5 filaments on immunostimulatory dsRNA molecules^11-13^. Hyperactive MDA5 alleles that exhibit enhanced dsRNA-activated IFN production are associated with higher risk of T1D, whereas hypomorphic alleles of MDA5 with reduced signaling activity correlate with protection from T1D^6-8^. An elevated type I IFN gene expression signature is detected in children with high genetic risk of diabetes prior to diabetes onset^14^, and patients with recent onset T1D exhibit higher expression of MDA5 in the pancreas^15^. In the NOD mouse, there are conflicting reports of the contribution of *Ifih1*/MDA5 to T1D onset. In one study, backcross of an *Ifih1* knockout allele from C57BL/6 mice into the NOD background completely protected from disease^16^. However, another study found that NOD mice engineered to have a small, in-frame deletion in the helicase domain of MDA5 were strongly protected from disease, but mice with a frameshift mutation resulting in a truncated MDA5 protein exhibited accelerated diabetes onset specifically in males^17^. The potential residual expression of the two CARD domains in the truncated MDA5 protein in these mice complicates interpretation of these findings, since the isolated CARDs can signal in a ligand-independent manner^18^. Finally, female NOD mice with a knockin of the *A946T* risk allele of *Ifih1* developed accelerated diabetes accompanied by higher expression of IFNs and IFN-stimulated genes^19^. Interestingly, onset of diabetes was not significantly changed in male *A946T/A946T* mice. Thus, the contribution of MDA5 to T1D in the NOD model remains incompletely defined.

We set out to explore the innate immune basis of T1D in the NOD mouse by applying lessons learned from another human disease called Aicardi-Goutieres Syndrome (AGS)^20^. AGS is an “interferonopathy” characterized by excessive antiviral responses to endogenous nucleic acids, and is caused by mutations in any of nine human genes that regulate intracellular nucleic acid detection^21,22^. Interestingly, rare activating mutations in the *IFIH1* gene cause a subset of AGS^23^, suggesting a potential mechanistic overlap with the genetics of human T1D. Moreover, AGS-causing mutations in the *ADAR* gene that encodes the ADAR1 RNA deaminase result in the accumulation of immunostimulatory RNAs that are specifically detected by MDA5^24^. We and others previously found that *Adar*-deficient mice, which are embryonic lethal, are rescued to birth by knockout of *Ifih1* (MDA5) *or Mavs*^25-27^. We then developed a knockin mouse model of a specific AGS point mutation in the *ADAR* gene, *Adar P195A. Adar*^*P195A/-*^ mice are born at normal Mendelian frequencies but develop severe disease with a median lifespan of ∼4 weeks^28^. These mice are completely rescued from disease when crossed to mice deficient for MDA5 (*Ifih1*), the type I interferon receptor (*Ifnar1*), the dsRNA-dependent kinase PKR (*Eif2ak2*), or Z DNA-binding protein 1 (*Zbp1*)^28,29^. We proposed a model in which ADAR1 mutation results in the accumulation of immunostimulatory RNAs that activate MDA5-mediated type I IFN production, which increases expression of the IFN-inducible RNA sensors PKR and ZBP1. PKR binds to immunostimulatory RNAs and phosphorylates eukaryotic initiation factor 2 alpha (EIF2α), which triggers the integrated stress response (ISR), resulting in global attenuation of protein synthesis and the upregulation of stress response genes^28^. Finally, ZBP1 activation, licensed by the ISR, drives inflammation in part through the induction of necroptotic cell death^29^. We found that treatment of *Adar*^*P195A/-*^ mice with the small molecule ISR inhibitor 2BAct rescued them from disease pathology and mortality, demonstrating the therapeutic potential of ISR inhibition in treating MDA5-mediated immune disease^28^. Importantly, NOD mice exhibit activation of the ISR in the pancreas^30^, and human T1D patients have altered ISR gene expression, including upregulation of *EIF2AK2* (PKR)^15^.

Given these potential genetic and mechanistic parallels between T1D and AGS, we used CRISPR technology to disrupt *Ifih1, Eif2ak2*, and *Zbp1* directly in the NOD background. We found that *Ifih1* heterozygosity resulted in delayed T1D onset specifically in male NOD mice, but complete *Ifih1* deficiency had no impact on disease. Moreover, loss of *Eifak2* or *Zbp1* had no influence on the onset or progression of T1D in either males or females. Finally, we found that treatment of NOD mice from gestation with 2BAct accelerated disease onset and increased diabetes incidence specifically in males, but had no effect on T1D in females. Our findings demonstrate that the MDA5-PKR-ZBP1 axis is not essential for disease in the NOD mouse model, and they show that *Ifih1* gene dosage and ISR activation contribute to sex-specific disease outcomes.

## Results

### MDA5 haploinsufficiency delays T1D disease onset specifically in male NOD mice

To determine how MDA5 affects T1D onset in NOD mice, we used CRISPR to target exon 1 of *Ifih1* in fertilized mouse oocytes (Figure 1A). Importantly, targeting of this first exon disrupts the CARD domains of MDA5, resulting in a null *Ifih1* allele. A founder mouse carrying a 315 bp deletion between nucleotides 293 and 609 and a 2 bp deletion between nucleotides 617 and 620, causing a premature stop codon (TAA) at AA 132, was used to establish a stable line of NOD.*Ifih1*^*-/-*^ mice (Figure 1B-C). Detection of the deletion allele was confirmed by PCR and DNA sequencing (Figure 1D-E). We intercrossed *Ifih1*^*+/-*^ NOD mice to generate cohorts of *Ifih1*^*+/+*^, *Ifih1*^*+/-*^ and *Ifih1*^*-/-*^ NOD mice. Pups born to NOD.*Ifih1*^*+/-*^ crosses were born at the expected Mendelian ratios (Figure 1F). We evaluated MDA5 protein expression in bone marrow-derived macrophages (BMDMs) derived from *Ifih1*^*+/+*^ (wild-type, WT), *Ifih1*^*+/-*^ and *Ifih1*^*-/-*^ NOD mice (Figure 1G). We found that MDA5 protein, which was expressed at low basal levels and was highly IFN-inducible, was reduced in *Ifih1*^*+/-*^ and completely absent from *Ifih1*^*-/-*^ cells (Figure 1G). Finally, we assessed spontaneous T1D onset in *Ifih1*^*+/+*^, *Ifih1*^*+/-*^ and *Ifih1*^*-/-*^ NOD mice. We found that male NOD mice heterozygous for MDA5 exhibited a significant delay and reduced incidence of T1D onset compared to WT male NOD mice (*P* < 0.05), whereas male *Ifih1*^*-/-*^ NOD mice developed T1D with similar kinetics to WT male NOD mice (Figure 1H). In contrast, female NOD mice developed T1D with identical onset and incidence regardless of *Ifih1* genotype (Figure 1I).

**Figure 1.**
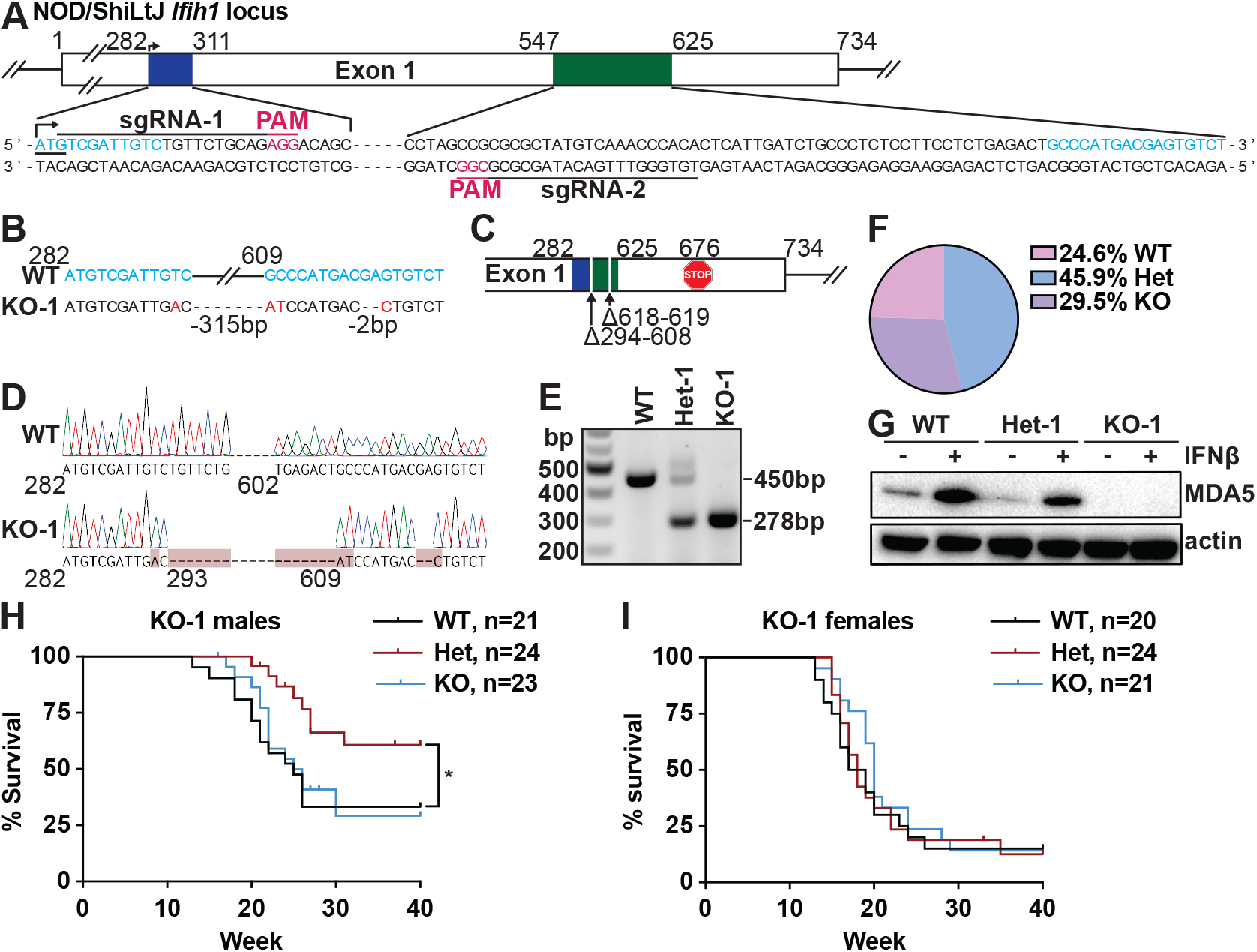
MDA5 haploinsufficiency reduces T1D disease onset in male NOD mice. (**A**) Schematic of the NOD/ShiLtJ *Ifih1* locus with CRISPR gRNA targeting sites. (**B**) DNA sequence of wild-type (WT) and *Ifih1* mutant alleles with SNPs in red. (**C**) Schematic of the *Ifih1* mutant allele. (**D**) Chromatogram of WT and *Ifih1* mutant alleles following PCR and Sanger sequencing. (**E**) Agarose gel electrophoresis analysis of PCR products from *Ifih1*^*+/+*^ (WT), *Ifih1*^*+/-*^ (Het) and *Ifih1*^*-/-*^ (KO) tissue. (**F**) Percentage of mice of the indicated genotype born from Het intercrosses. (**G**) Western blot for MDA5 on cell lysates from IFN-β-treated WT, Het and KO BMDMs. (**H** and **I**) Kaplan-Meier survival curves of diabetes incidence in WT, Het, and KO male (H) and female (I) mice. Analyzed by log-rank (Mantel-Cox) test; ^*^*p* < 0.05. Figures are representative of two (G) or at least three (D and E) independent experiments. (**H**) *n* = 21–24, (**I**) *n* = 20–24.

### Male NOD mice heterozygous for a “dominant negative” MDA5 allele are not protected from T1D onset

We established a second line of NOD.*Ifih1*-targeted mice using a founder mouse carrying a 268 bp deletion between nucleotides 296 and 565, causing a premature stop codon (TGA) at AA 118 (Figure 2A-C). We confirmed the presence of the deletion allele by PCR and DNA sequencing (Figure 2D), and pups born to NOD.*Ifih1*^*+/-*^ crosses in this line were again born at the expected Mendelian ratios (Figure 2E). However, and unlike the first NOD.*Ifih1* mouse line described in Figure 1, we found that MDA5 protein expression was undetectable in BMDMs from *Ifih1*^*+/-*^ mice (Figure 2F). Although the reasons for this complete loss of protein expression in cells from heterozygous mice are unclear, this suggests that the *Ifih1* mutant allele in this line may act as a dominant negative that precludes protein expression of the WT allele. Thus, at the level of protein expression, these *Ifih1*^*+/-*^ mice appear to be functional knockouts. In this line, we did not observe a protective effect of *Ifih1* heterozygosity in male mice (Figure 2G), and we again found that female mice had identical onset and incidence of diabetes regardless of *Ifih1* genotype (Figure 2H). Together, these findings in two independent lines of *Ifih1*-targeted NOD mice suggest that haploinsufficiency for *Ifih1*, accompanied by reduced MDA5 protein expression, has a modest protective effect specifically in male mice.

**Figure 2.**
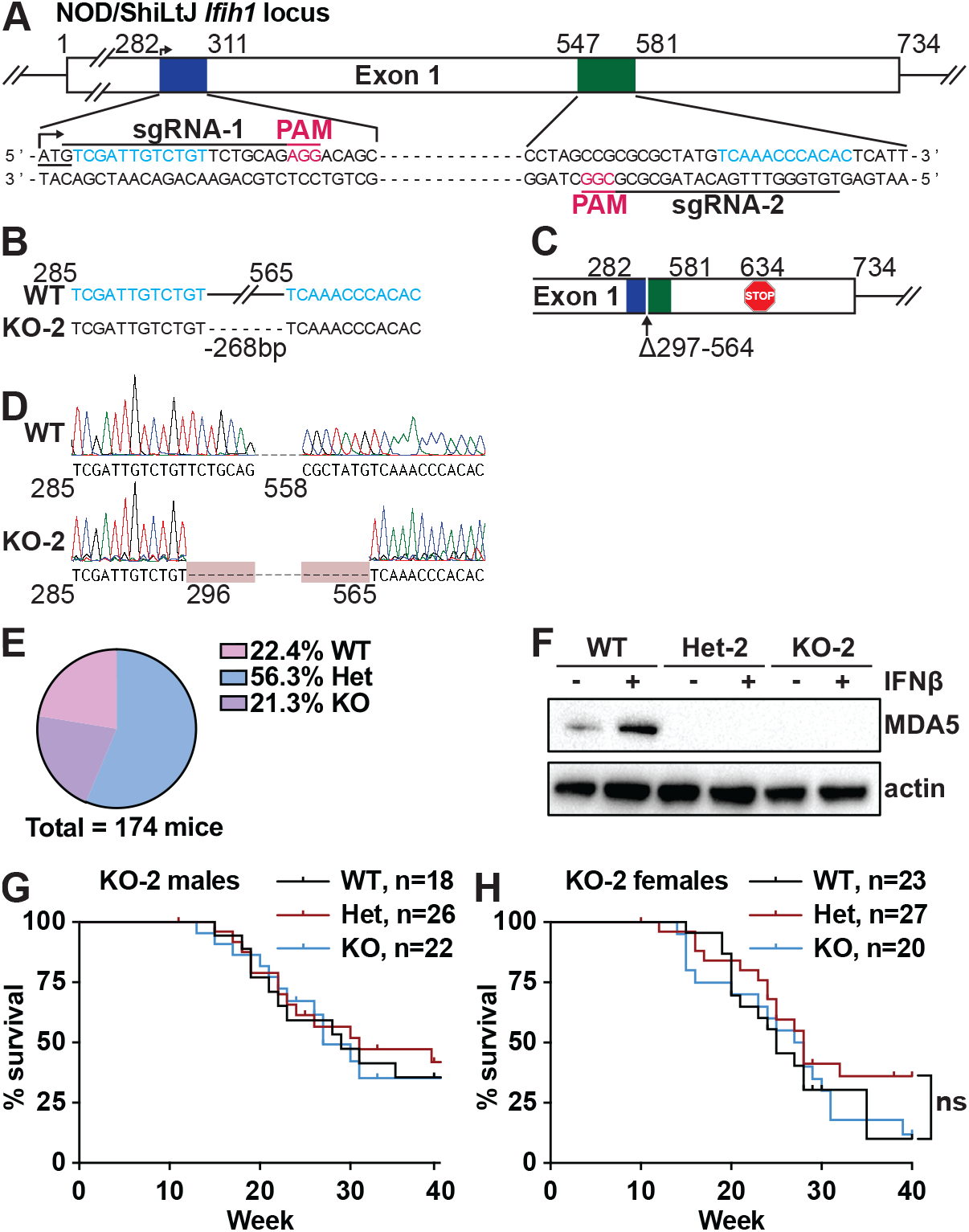
Male NOD mice heterozygous for a dominant negative MDA5 allele are not protected from T1D onset. (**A**) Schematic of the NOD/ShiLtJ *Ifih1* locus with CRISPR gRNA targeting sites. (**B**) DNA sequence of wild-type (WT) and *Ifih1* mutant alleles. (**C**) Schematic of the *Ifih1* mutant allele. (**D**) Chromatogram of WT and *Ifih1* mutant alleles following PCR and Sanger sequencing. (**E**) Percentage of mice of the indicated genotype born from Het intercrosses. (**F**) Western blot for MDA5 on cell lysates from IFN-β-treated WT, Het and KO BMDMs. (**G** and **H**) Kaplan-Meier survival curves of diabetes incidence in WT, Het, and KO male (H) and female (I) mice. Analyzed by log-rank (Mantel-Cox) test; ^*^*p* < 0.05. Figures are representative of two (F) or at least three (D) independent experiments. (**H**) *n* = 18– 22, (**I**) *n* = 20–27.

### Loss of PKR does not influence onset or incidence of T1D in NOD mice

To examine the effect of PKR on autoimmune diabetes, we used CRISPR to target Exon 2 of *Eif2ak2* in NOD mice (Figure 3A). Two founder mice carrying 1) a 423 bp deletion between nucleotides 4005 and 4429 or 2) a 39 bp deletion between nucleotides 4305 and 4345 and a 4 bp deletion between nucleotides 4424 and 4429, both with a loss of the canonical ATG start site, were used to establish two independent lines of NOD.*Eif2ak2*^*-/-*^ mice (Figure 3B-E). We confirmed the deletion alleles by PCR and DNA sequencing (Figure 3F-I), and we found that PKR protein expression was undetectable in BMDMs derived from both lines of NOD.*Eif2ak2*^*-/-*^ mice (Figure 3J-K). Pups born to NOD.*Eif2ak2*^*+/-*^ crosses were born at the expected Mendelian ratios for both lines (Figure 3L-M). We followed cohorts of both lines of NOD.*Eif2ak2* mice for 40 weeks, and we found that haploinsufficiency or loss of PKR did not significantly change the onset or incidence of diabetes in either male or female mice. (Figure 3N-Q).

**Figure 3.**
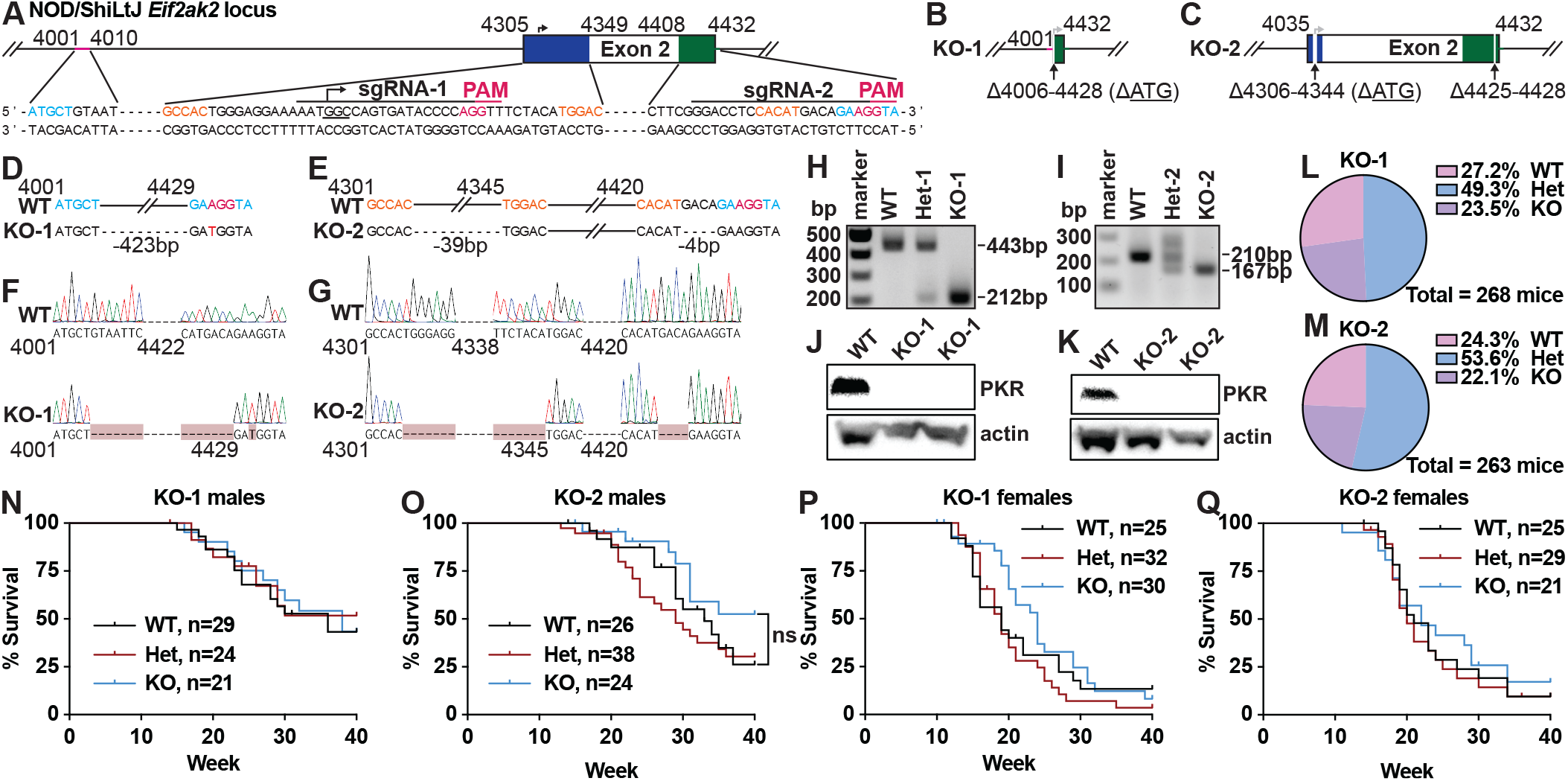
No difference in T1D onset following PKR targeting of NOD mice. (**A**) Schematic of the NOD/ShiLtJ *Eif2ak2* locus with CRISPR gRNA targeting sites. (**B** and **C**) Schematics of *Eif2ak2* mutant alleles KO-1 (B) and KO-2 (C). (**D** and **E**) DNA sequences of wild-type (WT) and *Eif2ak2* mutant alleles KO-1 (D) and KO-2 (E). (**F** and **G**) Chromatograms of WT and *Eif2ak2* mutant alleles KO-1 (F) and KO-2 (G). (**H** and **I**) Agarose gel electrophoresis analysis of PCR products from *Eif2ak2*^*+/+*^ (WT), *Eif2ak2*^*+/-*^ (Het) and *Eif2ak2*^*-/-*^ (KO) tissue for alleles KO-1 (H) and KO-2 (I). (**J** and **K**) Western blots for PKR on cell lysates from WT and KO-1 (J) and KO-2 (K) BMDMs. (**L** and **M**) Percentage of mice of the indicated genotype born from Het intercrosses in NOD.*Eif2ak2* lines KO-1 (L) and KO-2 (M). (**N**–**Q**) Kaplan-Meier survival curves of diabetes incidence in WT, Het, and KO male (N, O) and female (P, Q) mice from lines KO-1 (N, P) and KO-2 (O, Q). Analyzed by log-rank (Mantel-Cox) test. Figures represent data from two (J, K) or at least three (F–I) independent experiments. (**N**) *n* = 21–29, (**O**) *n* = 24–38, (**P**) *n* = 25–32, (**Q**) *n* = 21–29.

### ZBP1 does not contribute to diabetes in NOD mice

To investigate whether ZBP1 affects T1D onset in NOD mice, we used CRISPR to target Exon 2 of *Zbp1* (Figure 4A). Two founder mice carrying 1) a 124 bp deletion between nucleotides 1950 and 2075 causing a premature stop codon (TAG) at AA 68 or 2) a 130 bp deletion between nucleotides 1954 and 2085 causing a premature stop codon (TAA) at AA 95, were used to establish two lines of NOD.*Zbp1*^*-/-*^ mice (Figure 4B-E). We confirmed the alleles by both PCR and DNA sequencing (Figure 4F-I), and we found that both basal and IFN-inducible expression of ZBP1 protein was absent in BMDMs derived from NOD.*Zbp1*^*-/-*^ mice (Figure 4J). Pups born to NOD.*Zbp1*^*+/-*^ intercrosses were born at the expected Mendelian ratios for both lines (Figure 4K-L). We observed no differences in the sex-specific onset or incidence of T1D in either male or female *Zbp1*^*+/-*^ mice or *Zbp1*^*-/-*^ mice compared with WT NOD mice (Figure 4M-P).

**Figure 4.**
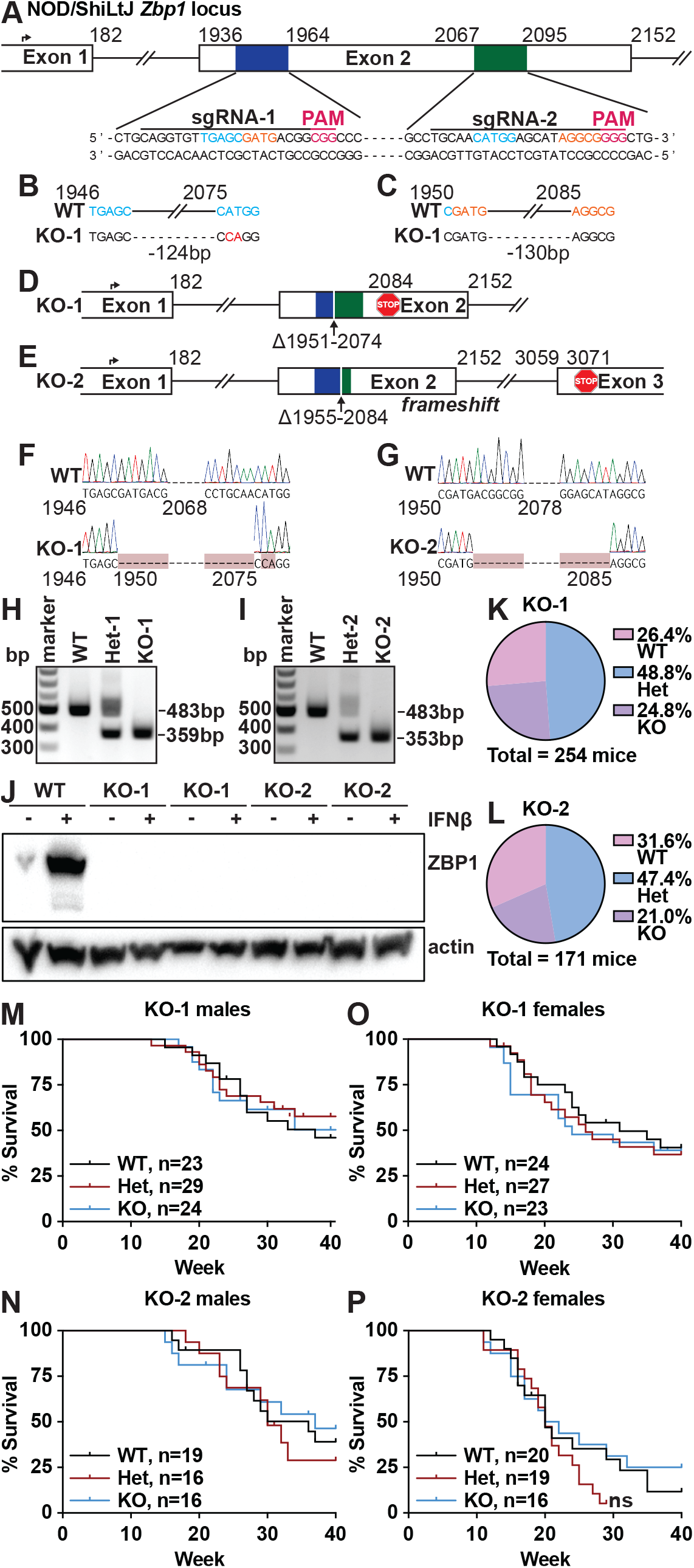
No difference in T1D onset following ZBP1 targeting of NOD mice. (**A**) Schematic of the NOD/ShiLtJ *Zbp1* locus with CRISPR gRNA targeting sites. (**B** and **C**) DNA sequences of wild-type (WT) and *Zbp1* mutant alleles KO-1 (B) and KO-2 (C) with SNPs in red. (**D** and **E**) Schematics of wild-type (WT) and *Zbp1* mutant alleles KO-1 (D) and KO-2 (E). (**F** and **G**) Chromatograms of WT and *Eif2ak2* mutant alleles KO-1 (F) and KO-2 (G). (**H** and **I**) Agarose gel electrophoresis analysis of PCR products from *Zbp1*^*+/+*^ (WT), *Zbp1*^*+/-*^ (Het) and *Zbp1*^*-/-*^ (KO) tissue for alleles KO-1 (H) and KO-2 (I). (**J**) Western blot for ZBP1 on cell lysates from IFN-β?-treated WT, KO-1, and KO-2 BMDMs. (**K** and **L**) Percentage of mice of the indicated genotype born from Het intercrosses in NOD.*Zbp1* lines KO-1 (K) and KO-2 (L). (**M**–**P**) Kaplan-Meier survival curvesof diabetes incidence in WT, Het, and KO male (M, N) and female (O, P) mice from lines KO-1 (M, O) and KO-2 (N, P). Analyzed by log-rank (Mantel-Cox) test. Figures represent data from two (J) or at least three (F–I) independent experiments. (**M**) *n* = 23–29, (**N**) *n* = 16–19, (**O**) *n* = 23–27, (**P**) *n* = 16–20.

### Pharmacological inhibition of the ISR worsens diabetes in male NOD mice

PKR is one of four eIF2α kinases that couple diverse cellular stresses to a program called the Integrated Stress Response (ISR), which restricts new protein synthesis and results in the induction of genes that can either restore homeostasis or cause cell death, depending on the strength and duration of the stress signal^31,32^. The eIF2 GTPase is responsible for the delivery of the initiator methionyl tRNA to the ribosome to commence mRNA translation at the AUG start codon. Phosphorylation of the α subunit of eIF2 on serine 51 prevents recycling of this eIF2 complex by the guanine nucleotide exchange factor (GEF) eIF2B, and it results in a reduction of most canonical mRNA translation initiation^33^. However, certain mRNAs that contain unusual arrangements of AUG start codons in their 5’ untranslated regions become specifically translated after eIF2α phosphorylation^34^. These include the transcription factor ATF4, which induces the expression of ISR-activated genes^35^. In addition to PKR, which detects dsRNA, the eIF2α kinase PERK responds to endoplasmic reticulum stress and triggers the ISR. Interestingly, the episodic production and secretion of insulin by pancreatic beta cells results in ER stress and PERK activation^36^. One recent study found that treatment of female NOD mice with a chemical inhibitor of PERK starting at 6 weeks of age delayed the onset and reduced the incidence of T1D^37^.

We previously found that long-term treatment of *Adar*^*P195A/-*^ mice with the orally bioavailable ISR inhibitor 2BAct, which binds to eIF2B and increases its activity^38^, prevented disease pathology and mortality^28^. We therefore hypothesized that 2BAct might ameliorate T1D in NOD mice. To test this, we placed NOD mice on control chow or chow containing 2BAct, starting with breeders and continuing after weaning of litters (Figure 5A). We monitored weights and T1D onset over 40 weeks of continuous treatment post-birth for both male and female NOD mice. We found that male mice treated with 2BAct lost significantly more weight than control-treated male NOD mice (Figure 5B). In contrast, 2BAct-treated female mice gained more weight compared to control-treated females (Figure 5C), suggesting sex-specific differences in response to ISR inhibition. 2BAct treatment had no effect on the onset or overall incidence of T1D in female NOD mice (Figure 5D). However, 2BAct treatment of male mice caused a significant acceleration in T1D onset, as well as an increase in the incidence of T1D (Figure 5D). This accelerated onset and increased incidence of 2BAct-treated male mice mirrored precisely the kinetics and penetrance of T1D in plain female NOD mice (Figure 5D), suggesting that the ISR may offer selective protection from T1D in male mice.

**Figure 5.**
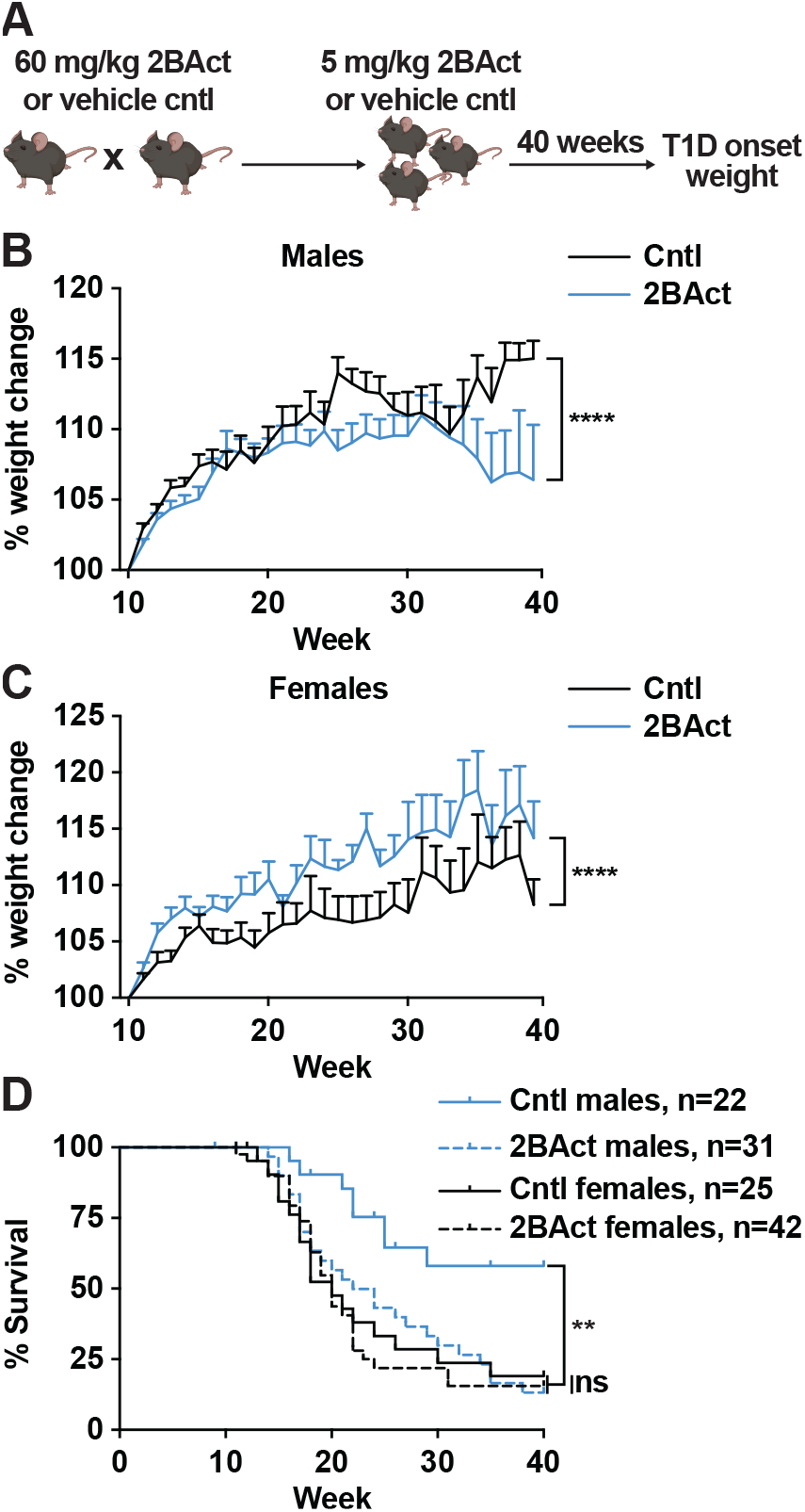
Inhibition of the integrated stress response accelerates T1D onset in male NOD mice. (**A**) Experimental design for *in vivo* 2BAct treatment of NOD/ShiLtJ mice. (**B** and **C**) ^%^ weight change from 10–40 weeks for male (B) and female (C) mice treated with 2BAct or control chow. (**D**) Kaplan-Meier survival curves of diabetes incidence in male and female mice treated with 2BAct or control chow. Analyzed by 2-way ANOVA (B, C) or log-rank (Mantel-Cox) test (D); ^**^*p* < 0.01, ^****^*p* < 0.0001. (**B)** *n* = 22-31, (**C**) *n* = 25-42, (**D**) *n* = 22–42.

## Discussion

In this study, we tested whether the genetic pathway and disease mechanisms that underlie MDA5-dependent disease in a mouse model of AGS apply to T1D in NOD mice, in which MDA5 has been strongly implicated as an important driver of pathology. To do this, we generated two lines each of NOD mice targeted for *Ifih1* (MDA5), *Eif2ak2* (PKR), and *Zbp1*, and we followed independent cohorts of all of six lines for 40 weeks to quantitate the progression of T1D. Overall, we did not observe significant changes in the onset or incidence of T1D in any of these homozygous knockout mice, demonstrating that the MDA5-PKR-ZBP1 axis is not responsible for disease development in the NOD model. This is in contrast to the complete rescue from pathology and mortality in the *Adar*^*P195A/-*^ mouse model of AGS on *Ifih1*^*-/-*^, *Eif2ak2*^*-/-*^ or *Zbp1*^*-/-*^ backgrounds^28,29^. However, we observed that haploinsufficiency for *Ifih1* modestly but significantly delayed T1D onset, but only in male NOD mice, whereas complete *Ifih1* deficiency had no influence on disease. This nonlinear gene dosage effect is unusual and suggests that the residual, reduced levels of MDA5 protein in *Ifih1*^*+/-*^ NOD mice may interact with other factors to offer this modest protection. Together with the previous reports characterizing MDA5 in NOD mice^16,17,19^, our findings suggest that the contributions of MDA5 to T1D are complex, sex-specific, subject to gene dosage effect, and perhaps different depending on whether the alleles are hyperactive risk alleles (which worsen disease in female NOD mice^19^), or hypomorphic expression alleles (which modestly protect male NOD mice). Additionally, our study focused on NOD mice housed in a specific pathogen free facility, in which infections that are thought to contribute to the onset of some cases of human T1D are not present. Such environmental perturbations might reveal a more pronounced influence of this pathway on disease.

Based on the proposed pathogenic role of the ISR in mouse and human T1D^36,37,39^, as well as the rescue of *Adar*^*P195A/-*^ mice from disease by treatment with the ISR inhibitor 2BAct, we tested whether 2BAct would modulate disease onset in NOD mice. Remarkably, we found that continuous 2BAct treatment had no protective effect in female NOD mice, and it accelerated the onset and worsened the incidence of disease in male NOD mice. The kinetics of T1D onset was identical between 2BAct-treated male NOD mice and control female NOD mice, which typically develop T1D faster and with higher penetrance than control male NOD mice. These findings suggest that the sex-specific differences in disease onset in the NOD model might be due in part to the differential protective role of the ISR in female versus male mice. Importantly, prior studies demonstrating protection from T1D through PERK inhibition^37^ or chemical amelioration of ER stress^39^ used female mice exclusively. In our studies, we found that PKR deficiency had no effect on T1D in either male or female NOD mice (Figure 3). Thus, we suggest that PKR is not the source of the ISR responsible for the male-specific exacerbation of disease caused by ISR inhibition. Instead, PERK is a likely source. More broadly, the ISR has emerged as a promising target for therapeutic inhibition, as exemplified by mouse models of vanishing white matter disease, AGS, and Pelizaeus-Merzbacher disease^28,38,40^. Whereas 2BAct is well tolerated in “normal” WT mouse models like C57BL/6^28,38^, our findings suggest that underlying conditions like the predisposition to T1D in NOD mice, as well as sex-specific differential responses to ISR inhibition, should be important considerations when deploying ISR-targeted therapeutics.

Together, our studies provide a stringent test of the contributions of MDA5, PKR, ZBP1, and the ISR to T1D in the NOD mouse model. Whereas all of these are essential for disease in the *Adar*^*P195A/-*^ mouse model of AGS, we show that development of T1D occurs independently of MDA5, PKR, and ZBP1, and that pharmacological ISR inhibition worsens disease in male NOD mice. These findings clarify important details about the origins and progression of T1D in the NOD mouse.

## Methods

### Mice

NOD/ShiLtJ (wild-type, strain 001976) mice were purchased from The Jackson Laboratory and bred in-house.*Ifih1*-targeted, *Eif2ak2*-targeted, and *Zbp1*-targeted NOD/ShiLtJ mice were generated by microinjecting oocytes from wild-type NOD/ShiLtJ mice with Cas9 complexed with two gRNAs targeting *Ifih1* exon 1 (TGTGGGTTTGACATAGCGCG and GTCGATTGTCTGTTCTGCAG), *Eif2ak2* exon 2 (AAAATGGCCAGTGATACCCC and CGGGACCTCCACATGACAGA), or *Zbp1* exon 2 (CAGGTGTTGAGCGATGACGG and TGCAACATGGAGCATAGGCG). Oocytes were implanted into pseudopregnant CD-1 females via oviduct transfer. Founder pups were genotyped by PCR and Sanger sequencing using the following sequencing primers:

NOD.*Ifih1* FWD (common) AGCTCCCTGAGGGTGAACGTCC NOD.*Ifih1* REV (WT) AGGTGCTCAGCAGCAGTTCTGC

NOD.*Ifih1* REV (KO-1) GGAGACAGGTCATGGATGTCAATCGACATCG NOD.*Ifih1* REV (KO-2) TGAGTGTGGGTTTGAACAGACAATCG NOD.*Eif2ak2* FWD (common) TGTGTTTGCCCCAACAAAGCAC

NOD.*Eif2ak2* REV (WT) TCTGGTGTTTCCAACCCACCAC NOD.*Eif2ak2* REV (KO-1) GGCAGCCTACCTTCGGCATTTA NOD.*Eif2ak2* FWD (KO-2) GTAGGCCCCTCTTCCTTTGCAG NOD.*Eif2ak2* REV (KO-2) TTGCTGACTGAGTCACCTCCCT NOD.*Zbp1* FWD TACCATTTGGCCACCCAGAC

NOD.*Zbp1* REV TCGTGGCTCTTAATGTGCGTGT

Founder pups were backcrossed to WT NOD/ShiLtJ mice for two generations to generate stable lines of heterozygous NOD.*Ifih1*^*+/-*^, NOD.*Eif2ak2*^*+/-*^, and NOD.*Zbp1*^*+/-*^ mice. Heterozygous mutant mice were intercrossed to generate littermate-controlled cohorts of WT, heterozygous, and knockout animals. Blood glucose levels were monitored weekly from 10-40 weeks of age using the AimStrip Plus Blood Glucose Testing System (Germaine Laboratories). Diabetes incidence was confirmed by 2 consecutive blood glucose readings ≥ 300 mg/dL. All mice were maintained in a specific pathogen-free (SPF) barrier facility at the University of Washington, and all experiments were done in accordance with the Institutional Animal Care and Use Committee guidelines of the University of Washington.

### Cell culture

Bone marrow was obtained from WT NOD, NOD.*Ifih1*^*+/-*^, NOD.*Ifih1*^*-/-*^, NOD.*Eif2ak2*^*-/-*^, and NOD.*Zbp1*^*-/-*^ mice and cultured in RPMI supplemented with 10% fetal calf serum, 2 mM L-Glutamine, 100 U/mL penicillin, 100 U/mL streptomycin, 1mM sodium pyruvate, 10 mM HEPES, and 10% supernatant from 3T3–M-CSF cells for 6 d with feeding on day 3 to generate bone marrow-derived macrophages (BMDMs). 2.7x10^5^ BMDMs were plated in 24-well plates and rested overnight, then stimulated with 100 U/mL recombinant murine IFN-β (R&D Systems, 12405-1) or equivalent volume of water for 24 hours. Cells were washed with PBS, lysed in RIPA buffer containing 1x protease inhibitor (ThermoFisher Scientific, A32953) for 15 min on ice, and centrifuged at 15,000 RPM for 15 min at 4C. Protein-containing supernatants were harvested and frozen at -80C for downstream western blot analysis.

### Western blot

Protein samples in 1x SDS sample buffer + β-mercaptoethanol were analyzed by SDS-PAGE using precast 4-12% polyacrylamide gels (ThermoFisher Scientific, NW04127BOX) and mAbs to MDA5 (Cell Signaling Technology, 5321S), PKR (Santa Cruz Biotechnology, sc-6282), or ZBP1 (AdipoGen, AG-20B-0010-C100) and HRP-conjugated anti-rabbit or anti-mouse secondary antibodies. Blots were developed with Pierce™ ECL 2 Western Blotting Substrate (ThermoFisher Scientific, PI80196x3) and a ChemiDoc MP system from Bio-Rad Laboratories.

### 2BAct treatments

Breeding pairs of WT NOD/ShiLtJ mice were placed on Teklad 2014 chow formulated to achieve a 2BAct concentration of 60 mg/kg. Upon weaning, pups were placed on chow formulated to achieve a 2BAct concentration of 5 mg/kg. The placebo diet was Teklad 2014 without added compound. 2BAct and placebo chow was contract manufactured with Envigo. Pups were maintained on 2BAct until terminal endpoint or 40 weeks of age. Pups were weighed weekly starting at 10 weeks and diabetes incidence was monitored as described above.

## Statistical analysis

Analysis of statistical significance was performed using GraphPad Prism 11 (GraphPad, La Jolla, CA).

## Acknowledgements

We thank Neal Mausolf for mouse colony management, Priti Singh and the Fred Hutchinson Cancer Research Center Preclinical Modeling Core for expertise in the generation of CRISPR-targeted mice, Marion Pepper for key reagents, Rebecca Hull-Meichle, Sakeneh Zraika, Elizabeth Giering and Breanne Barrow at the University of Washington Diabetes Research Center and Veterans Affairs Puget Sound Health Care System and Brian Johnson and Brendy Fountaine at the University of Washington Histology and Imaging Core for technical assistance, and all members of the Stetson lab for helpful discussions. This work was supported by NIH R01 AI084914 (D.B.S.) Erik Van Dis is a Robert Black Fellow of the Damon Runyon Cancer Research Foundation (DRG-2483-23).

## Author contributions

Conceptualization: Erik Van Dis, Daniel B. Stetson Formal analysis: Erik Van Dis

Funding acquisition: Erik Van Dis, Daniel B. Stetson

Investigation: Erik Van Dis, Alessandra T. DeGidio, Lara Yao, Damion Winship Methodology: Erik Van Dis, Daniel B. Stetson

Resources: Carmela Sidrauski, Jacquelyn A. Gorman Supervision: Daniel B. Stetson

Writing – original draft: Erik Van Dis, Daniel B. Stetson

Writing – review & editing: Erik Van Dis, Alessandra T. DeGidio, Daniel B. Stetson

## Declaration of Interests

C.S. is an employee of Calico Life Sciences and is listed as an inventor on a patent application WO2017193063 describing 2BAct.

